# Establishment of a Systemic Lupus Erythematosus Mouse Model in Humanized CD3/CD19 Mice and Evaluation of the Efficacy of a Human Bispecific Antibody

**DOI:** 10.1101/2025.08.21.668572

**Authors:** Yaqi Du, Lijun Zhou, Jiaxuan Shao, Yanling Wang, Menghan Cui, Shuang Li, Yu Su, Lei Ci, Ruilin Sun

**Author notes:** **Correspondence should be addressed to** Yaqi Du (Lead contact,). **Conflict of interest:** The authors declare no potential conflicts of interest.

## Abstract

Systemic Lupus Erythematosus (SLE) is a complex autoimmune disease affecting multiple organ systems and is considered one of the most severe autoimmune diseases worldwide, significantly impacting women’s health. B cells play a crucial role in disease pathogenesis by becoming abnormally activated and producing autoantibodies that attack the body’s normal tissues, leading to inflammation and organ damage. Therefore, targeting the human B cell receptor has emerged as a promising strategy for therapeutic intervention. In this study, we established an SLE animal model using double-humanized CD3/CD19 mice, in which human CD3 and CD19 genes replaced their murine counterparts. The phenotype was confirmed by ELISA and flow cytometry. hCD3/hCD19 mice induced by a combination of pristine x 2, imiquimod (IMQ) and nephrectomy exhibited several key features of SLE, including anti-dsDNA antibody production, proteinuria, increased immune cell populations, and kidney lesions. Furthermore, treatment of these humanized mice with anti-human CD3/CD19 bispecific antibodies significantly inhibited these pathological indicators. Thus, pristine x 2, IMQ and nephrectomy induced hCD3/hCD19 mice provide a validated preclinical model for investigating novel therapeutic agents targeting B cells in SLE.

**Graphical abstract:** This study explores the effects of different modeling strategies on agent induction, with a particular emphasis on the link between the induced agents and their corresponding clinical manifestations. Through an extensive literature review and experimental validation, a novel approach for inducing systemic lupus erythematosus (SLE) has been established. By employing gene editing techniques, mouse CD3 and CD19 antibody recognition sites were substituted with their human equivalents. This humanized mouse model enables the direct assessment of humanized bispecific antibodies in the context of SLE induction.

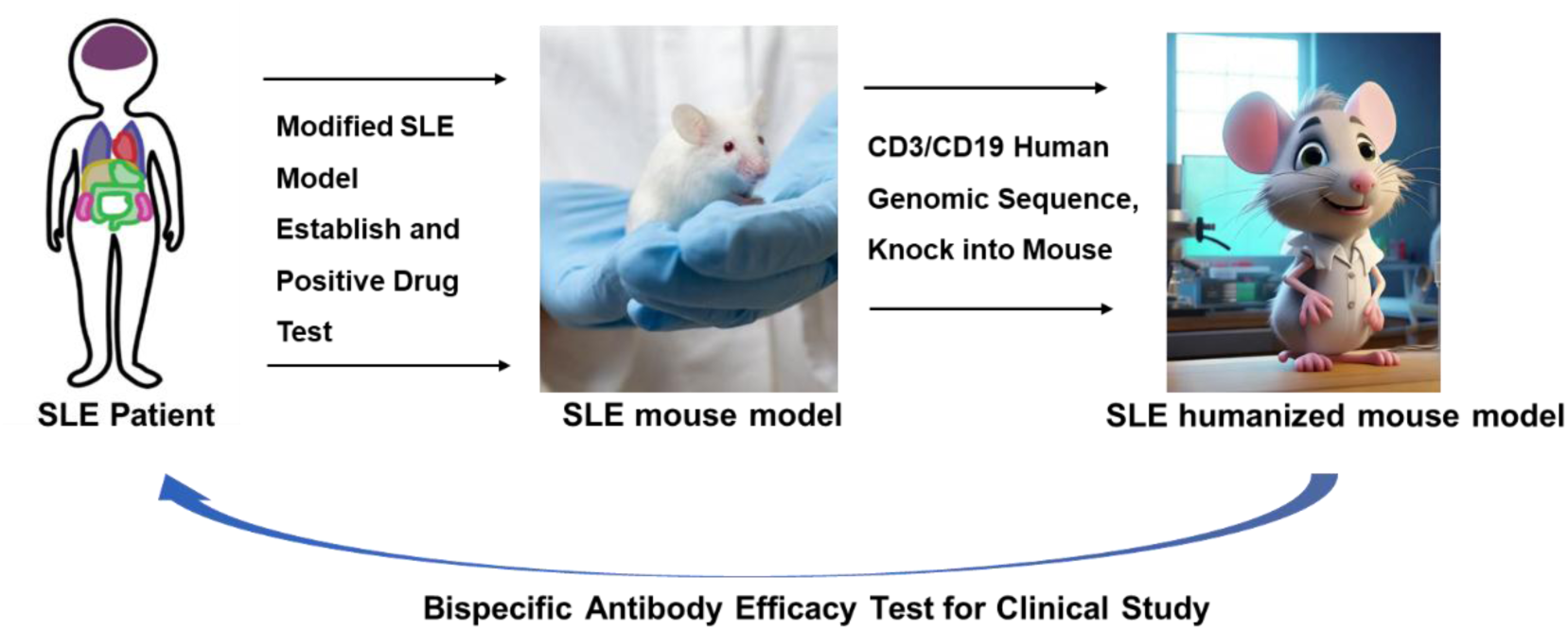

## 1 INTRODUCTION

Systemic lupus erythematosus (SLE) is a chronic, relapsing, and inflammatory autoimmune disorder that affects multiple organ systems, particularly the skin, joints, kidneys, and serous membranes^1^. Globally, it impacts approximately 3 million individuals, with women comprising over 90% of affected cases^1,2^. The disease primarily strikes women of childbearing age (20–40 years), severely impairing their quality of life and fertility^3^. Lupus nephritis (LN) represents one of the most serious complications associated with SLE^4^.

The pathogenesis of SLE is multifactorial, involving genetic predisposition, environmental triggers, and immune system dysregulation^5^. Aberrant expansion and activation of autoreactive B cells are critical in driving disease progression^6^. B cells contribute not only to antibody production but also to antigen presentation, co-stimulation, and immune modulation. Dysfunctional B cell differentiation leads to the generation of pathogenic autoantibodies, such as anti-double-stranded DNA (anti-dsDNA), anti-nuclear antibodies (ANA), and anti-Smith (anti-Sm) antibodies^7^. Elevated serum levels of anti-dsDNA antibodies serve as a key biomarker for SLE diagnosis and reflect underlying B cell-mediated autoimmunity^8^. The formation of immune complexes by these autoantibodies can activate the complement cascade, contributing to tissue injury and renal impairment^9,10^. Renal function is commonly monitored through urinary albumin measurements and urine albumin-to-creatinine ratio (UACR) calculations^11^. Serum anti-dsDNA titers, UACR levels, and kidney immunopathology, including IgG deposition, are important indicators of disease severity and progression^8,12,13^.

CD19, expressed throughout most stages of B cell development from precursor to plasma cells, plays a central role in B cell receptor (BCR) signaling by recruiting downstream kinases essential for activation and differentiation^14^. Similarly, the T-cell receptor (TCR)/CD3 complex, composed of TCR-α/β chains and six CD3 subunits (CD3δε, CD3γε, and CD3ζζ dimers), is vital for antigen recognition, signal transduction, and T cell development. However, the extracellular domains of human and mouse CD3 proteins exhibit limited homology (47% for CD3ε, 57% for CD3δ, and 60% for CD3γ), hindering cross-species therapeutic efficacy. As a result, agents targeting human CD3 cannot effectively stimulate mouse T cells via their endogenous CD3 complex. To overcome this limitation, fully humanized CD3EDG mice, engineered to express human CD3 domains, have been developed for evaluating human-specific CD3 bispecific antibodies in vivo.

Conventional SLE therapies primarily include immune-suppressants such as glucocorticoids, cyclophosphamide, and mycophenolate mofetil, alongside antimalarial agents like hydroxychloroquine^15^. However, prolonged use of these treatments is associated with significant adverse effects: steroid therapy can cause infections, hypertension, hyperglycemia, obesity, avascular necrosis, and neuropsychiatric disorders^16^; cyclophosphamide poses risks to gonadal function^17^; Mycophenolate mofetil increases susceptibility to infections^18^, and hydroxychloroquine may induce ocular toxicity, including retinopathy^19^.

In recent decades, biologic therapies targeting specific immune pathways with reduced toxicity profiles have emerged. Belimumab, the first biologic approved for SLE, inhibits B cell survival by neutralizing BAFF^20^. Telitacicept, a recombinant fusion protein, competitively blocks BAFF and APRIL cytokine activity ^21^, while monoclonal antibodies like anifrolumab target IFN pathways^22^. Despite these advances, the clinical efficacy of biologics remains suboptimal, highlighting the ongoing need for safe, durable therapeutic options. Consequently, new strategies have focused on targeting B cell surface markers such as CD20, CD19, and BCMA to enhance specificity and minimize off-target effects^23^. CD19-targeted therapies, including CAR-T cell therapy, have demonstrated promising results^24–26^; however, CAR-T approaches also present challenges, such as risks associated with viral vector integration, prolonged T cell persistence, complex manufacturing processes, high costs, and preconditioning regimens that may cause organ toxicity and infertility. In severe, refractory autoimmune diseases, bispecific T cell engagers (e.g., anti-CD3/anti-CD19 antibodies) offer an attractive alternative^27^.

Animal models have been instrumental in understanding SLE pathogenesis and evaluating therapeutics. Classical spontaneous models like NZB/W and MRL/lpr mice have been widely used since the 1960s^28–30^. However, as biologics and cell therapies increasingly target human-specific receptors and cytokines, traditional models face limitations^31^. Pristane-induced SLE models, though commonly used, require long induction periods and produce relatively mild disease manifestations^32^. Toll-like receptor 7 (TLR7), a key innate immune sensor, is essential for extrafollicular B cell activation and germinal center formation, both critical for autoantibody production^33^. Activation of TLR7 by imiquimod (IMQ) enhances IFN-γ production, promoting robust activation of T cells, B cells, macrophages, and dendritic cells. Co-stimulation through TLR7 thus provides a strategy to augment pristane-induced disease severity^34,35^.

To address these limitations, we developed a novel humanized mouse model, replacing murine CD3 and CD19 genes with their human counterparts, and optimized the induction protocol by combining pristine x2, IMQ treatment and nephrectomy. In this study, we demonstrate that this double-humanized model supports the evaluation of anti-human CD3/CD19 bispecific antibody efficacy in vivo. This platform provides a valuable tool for the preclinical development of next-generation bispecific therapeutics targeting B cells in SLE^36^.

## 2 METHODS

### 2.1 Model Induction Procedure

The modified SLE model based on BALB/C background mice was induced by first performing a nephrectomy. One-week post-surgery, 500 µL of pristane (Sigma, USA) was administered to the mouse via intraperitoneal injection. Subsequently, IMQ (3C Health, USA) was applied daily to the right ear of the mouse. On the 14th day, an additional 500 µL of pristane was administered intraperitoneally. The induction phase of the model is designed to span a total duration of two months.

### 2.2 flow-cytometry detection

The immunocyte counts in peripheral blood, spleen, and lymph nodes were detected using flow cytometry (Beckman Coulter CytoFLEX, USA). The spleen was processed with red blood cell lysis solution (Biolegend, USA) for grinding, and 100 µL/1 mL of the grinding solution was extracted for subsequent staining. Lymph nodes were treated with stain buffer (BD Biosciences, USA) and subjected to grinding before proceeding with staining. Peripheral blood was centrifuged and then incubated with red blood cell lysis solution. The solution was then incubated for 10 minutes in the dark at 4 °C with a mixture of live/dead stains (Biolegend, USA) and anti-mCD16/32 (Biolegend, USA) to block nonspecific binding. After an additional 30-minute incubation in the dark at 4 °C with a cocktail of appropriately labeled fluorescent monoclonal antibodies, the cells were washed with PBS and resuspended in FACS staining buffer (PBS with 2% FBS). The antibodies used against mouse cell-surface molecules are as follows (purchased from Biolegend, USA unless otherwise indicated) (targeting mouse antigens unless otherwise indicated): anti-CD45-Percp/CY5.5 (clone 30-F11), anti-CD3-AF700 (clone 17A2), anti-mouse CD8-APC (clone 53-6.7), anti-CD4-BV605 (clone RM4-4), anti-CD19-APC (clone 1D3), anti-CD138-PE/Cyanine7 (clone 281-2), anti-CD11b-BV650 (clone M1/70), anti-CD11c-FITC (clone N418), anti-MHCII-BV421 (clone M5/114.15.2). For intracellular cytokine staining, cells were fixed and permeabilized using the Zombie NIR™ Fixable Viability Kit (BioLegend, USA), and then stained with anti-Foxp-PE (clone MF-14). Following this, the cells were analyzed by multicolor flow cytometry on the Beckman Coulter CytoFLEX and the data were analyzed using FlowJo 10 software.

### 2.3 Elisa detection

Blood samples were drawn from the retrobulbar venous plexus. Serum was then separated by centrifugation, and the anti-double-stranded DNA (anti-dsDNA) antibody titer was measured using the Anti-dsDNA IgG Antibody ELISA Assay Kit (Chondrex, USA). Mouse IgG was measured using the IgG (Total) Mouse ELISA Kit (Thermo, USA), and mouse IgM was measured using the Mouse IgM ELISA Kit (abclonal, China).

### 2.4 Histological and IHC detection

Kidney and salivary gland tissues were prepared for histological analysis by fixation in 10% buffered formalin, followed by routine paraffin embedding. Sections of 5 μm were cut and stained with hematoxylin and eosin (H&E) or subjected to IgG immunohistochemistry (IHC) using antibodies from Southern Biotech (USA).Using Halo software to analyze the percentage of IgG-positive areas in the renal cortex, and finally, these tissues were scored using the revised ISN/RPS lupus nephritis scoring criteria to assess the increase in intracapillary cells, neutrophils/nuclear fragmentation, fibrinoid necrosis, hyaline deposits, cellular/fibro cellular crescents, interstitial nephritis (activity index); glomerulosclerosis, fibrous crescents, tubular atrophy, and interstitial fibrosis (chronicity index); and the salivary glands were scored according to the degree of inflammatory cell infiltration.

### 2.5 CFSE-dilution detection

CD3+ T cells were isolated in the splenocytes from WT BALB/c and hCD3/hCD19 mice. Isolated T cells were labeled with µCFSE dye and stimulated with anti-mCD3 (5 µg/mL) or anti-hCD3 (5 µg/mL) plus soluble anti-mCD28 (50 µg/mL) for in vitro culture for 72 hr. T cell proliferation was analyzed by flow cytometry for CFSE dilution.

### 2.6 UACR detection

The detection of UACR is carried out as follows: 24-hour urine samples from mice are collected using metabolic cages (Glo-bio, China), and then analyzed using an automatic biochemical analysis system (Hitachi, Japan). The detection items include urinary albumin (Medical system-bio, China), creatinine (Fuji, Japan), and the ratio of albumin to creatinine, UACR.

### 2.7 Data analysis

Results are presented as the mean ± SEM. Statistical analysis was performed using SPSS 19, and GraphPad Prism 6 was used for data visualization. Comparisons were made using Student’s t-test and one-way ANOVA. A p-value of < 0.05 was considered statistically significant. Specific p-value notations are as follows: * *p* < 0.05, ** *p* < 0.01, *** *p* < 0.001, and **** *p* < 0.0001.

## 3 RESULTS

### 3.1 A modified method to induced SLE mouse model in Balb/c mice

In this study, we established multiple induction protocols to compare differences in disease modeling (Fig. 1A). Across all induction groups, mouse body weights remained within a safe range, and the animals exhibited good tolerance to the positive control treatment with cyclophosphamide (Fig. 1B). Induction led to a significant increase in peripheral blood inflammatory cells, particularly under the combined pristane and IMQ stimulation protocol. Cyclophosphamide administration effectively reduced the elevated inflammatory cell counts (Fig. 1C).

**Fig. 1.**
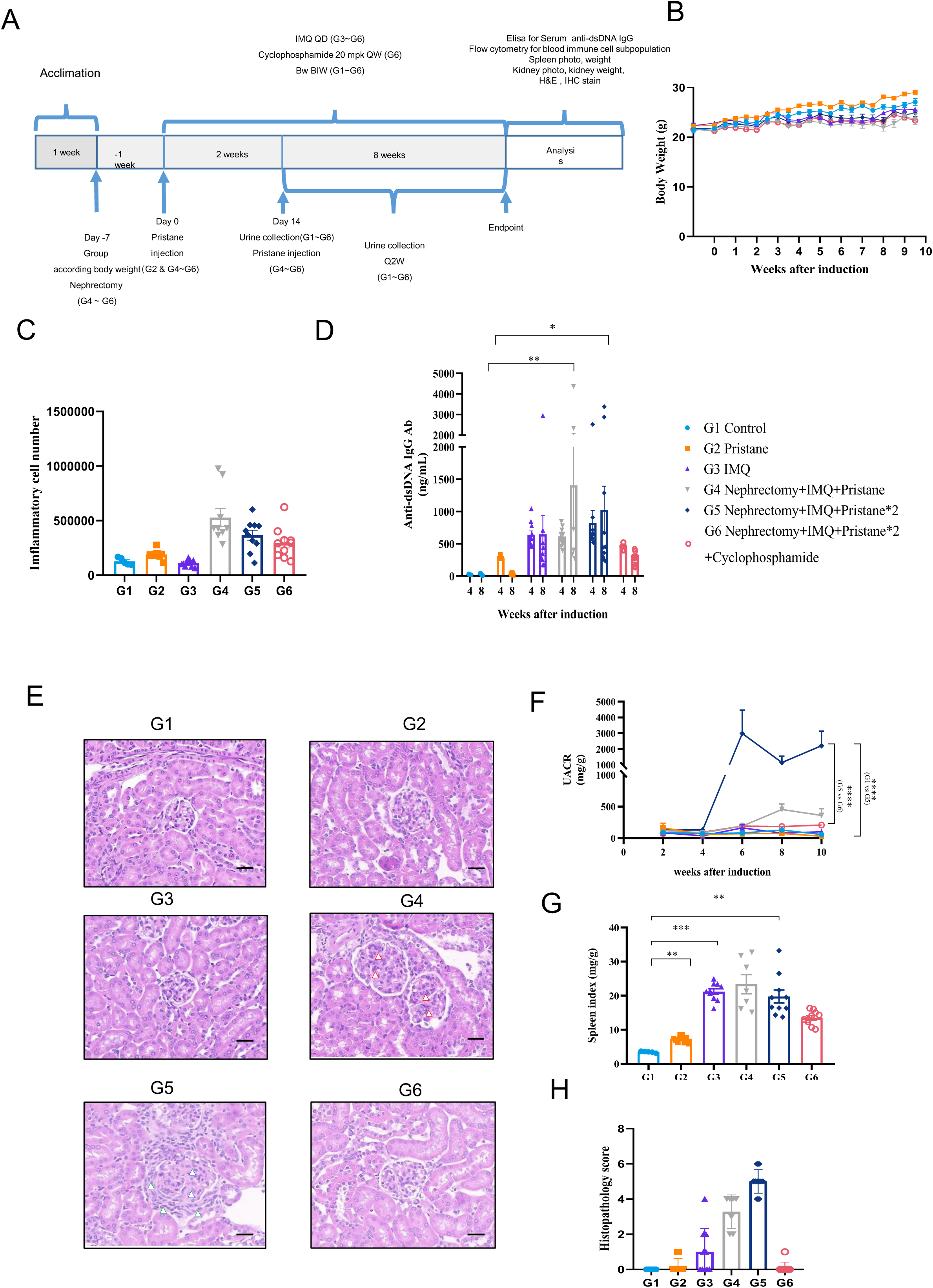
A modified approach for inducing an SLE mouse model in Balb/c mice. (A) Schematic illustration of the SLE model establishment. G1: untreated control group; G2: mice receiving a single 500 µL Pristane injection on Day 0; G3: mice subjected to daily IMQ application; G4: mice undergoing unilateral nephrectomy followed by one 500 µL Pristane injection and daily IMQ treatment (combined induction); G5: mice undergoing unilateral nephrectomy, two separate 500 µL Pristane injections, and daily IMQ application (enhanced combined induction); G6: mice treated according to G5 procedures, with additional cyclophosphamide therapy. (B) Body weight changes over time across all groups. (C) Quantification of inflammatory cells in 100 μL of peripheral blood at week 10. (D) Serum anti-dsDNA IgG levels in each group measured at weeks 4 and 8. (E) Representative H&E-stained kidney sections from each group at the experimental endpoint. (F) Urinary albumin-to-creatinine ratio (UACR) from 24-hour urine samples collected throughout the study. (G) Spleen index (spleen weight/body weight ratio) measured at the endpoint. (H) Quantitative scoring of kidney lesions across groups. Histological sections were visualized at 40× magnification; scale bar = 100 μm.

Flow cytometric analysis of peripheral blood immune cell subsets revealed a predominant expansion of myeloid cells and a concomitant reduction in lymphoid cells following disease induction (Supplemental Fig. 1A–1F). Serum levels of anti-double-stranded DNA (anti-dsDNA) antibodies were notably elevated at both 4- and 8-weeks post-induction, with the combined pristane and IMQ group showing the most pronounced increases; treatment with cyclophosphamide significantly suppressed anti-dsDNA antibody production (Fig. 1D).

Histopathological examination of renal tissues revealed significant kidney damage post-induction. Notably, mice subjected to unilateral nephrectomy, IMQ application, and double pristane injection exhibited the most severe pathological changes, characterized by glomerular mesangial hyperplasia and sclerosis, both of which were alleviated by cyclophosphamide treatment (Fig. 1E, 1H). Similar trends were observed in renal IgG deposition and salivary gland pathology (Supplemental Fig. 1I–1L).

In addition, model mice developed notable proteinuria, accompanied by a significant rise in urine albumin-to-creatinine ratio (UACR), both of which were markedly reduced by cyclophosphamide therapy (Fig. 1F; Supplemental Fig. 1G–1H). Splenomegaly was also observed in the model group, and treatment with cyclophosphamide effectively mitigated spleen enlargement (Fig. 1G).

### 3.2 T Cell and B Cell Functionality in hCD3/hCD19 Mice

The murine genes encoding CD3E, CD3D, and CD3G were substituted with their human counterparts (hCD3E, hCD3D, and hCD3G), and the extracellular domain of murine CD19 was replaced with human CD19 using gene-editing techniques (Supplemental Fig. 2A–2B). Following selective breeding, homozygous hCD3/hCD19 mice were successfully generated.

**Fig. 2.**
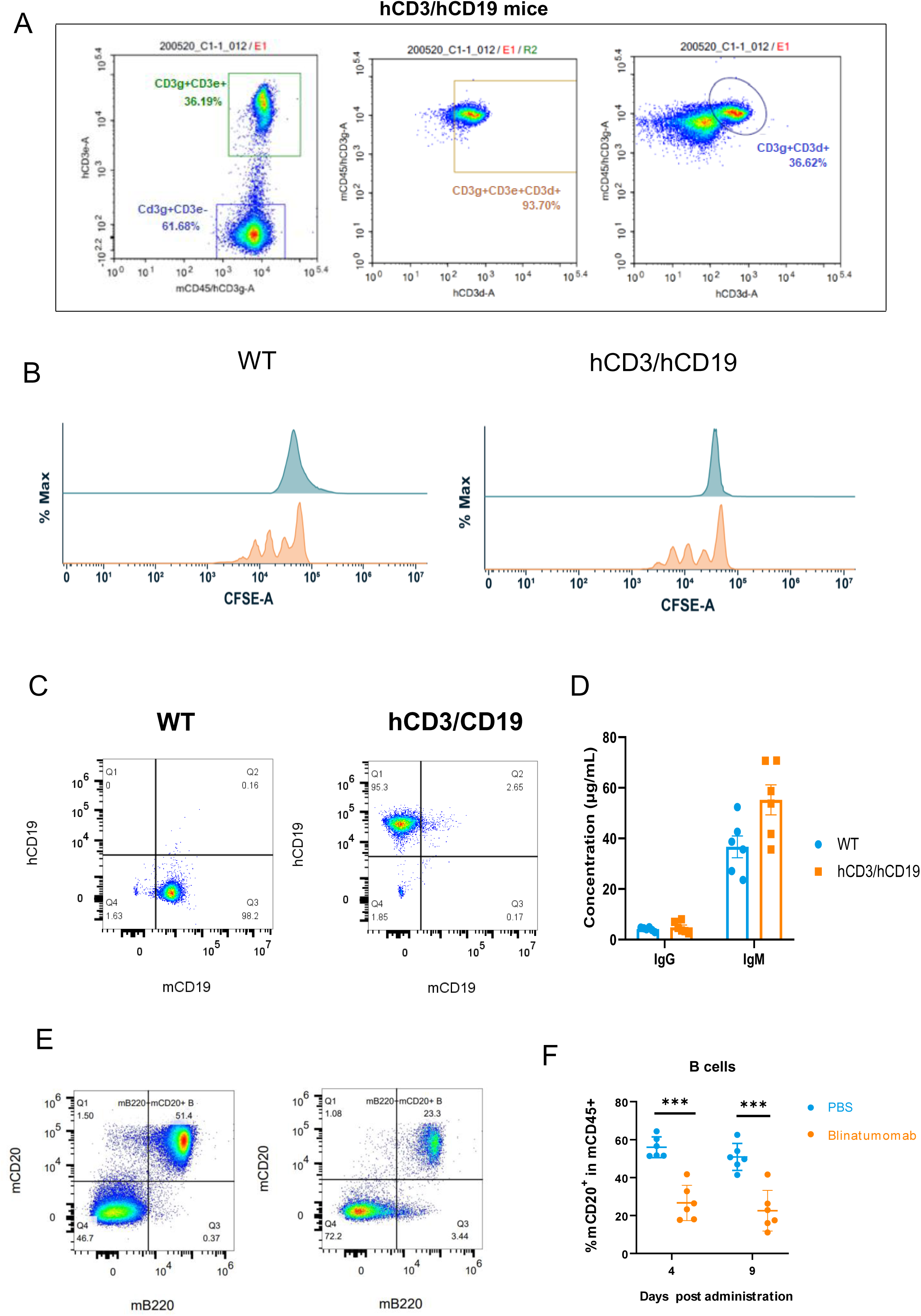
Functional assessment of T and B cells in hCD3/hCD19 mice. (A) Flow cytometry analysis of human CD3E, CD3D, and CD3G surface expression on splenocytes from hCD3/hCD19 mice. (B) Evaluation of TCR signaling following in vitro activation with CD3/CD28 antibodies. (C) Detection of human CD19 surface expression on splenic B cells. (D) Serum levels of IgG and IgM in hCD3/hCD19 mice. (E) Representative flow cytometry plot highlighting B cell populations. (F) Proportion of B cells among inflammatory cells.

Flow cytometric analysis confirmed the surface expression of hCD3E, hCD3D, and hCD3G proteins on splenic immune cells from hCD3/hCD19 mice (Fig. 2A). Upon stimulation with anti-CD3 and anti-CD28 antibodies, T cell proliferation in hCD3/hCD19 mice was comparable to that observed in wild-type controls (Fig. 2B). Similarly, the expression of hCD19 on splenic B cells was readily detected (Fig. 2C), and serum levels of immunoglobulins (IgG and IgM) showed no significant differences between hCD3/hCD19 mice and wild-type mice (Fig. 2D).

Following treatment with blinatumomab, a significant reduction in peripheral B cell numbers was observed in hCD3/hCD19 mice (Fig. 2E and 2F). Concurrently, an increase in the proportion of cytotoxic T cells and a decrease in the proportion of helper T cells were noted, while the overall T cell population remained largely unchanged (Supplemental Fig. 2C–2F).

### 3.3 Immune Cell Subpopulation Analysis in Different Immune Tissues

Flow cytometric analysis was performed to characterize immune cell subsets, including myeloid cells, B cells, T cells, helper T cells, cytotoxic T cells, and NK cells, in the peripheral blood, spleen, and lymph nodes of hCD3/hCD19 mice. The distribution of these immune cell populations showed no significant differences compared with those observed in wild-type mice (Fig. 3A–3F).

**Fig. 3.**
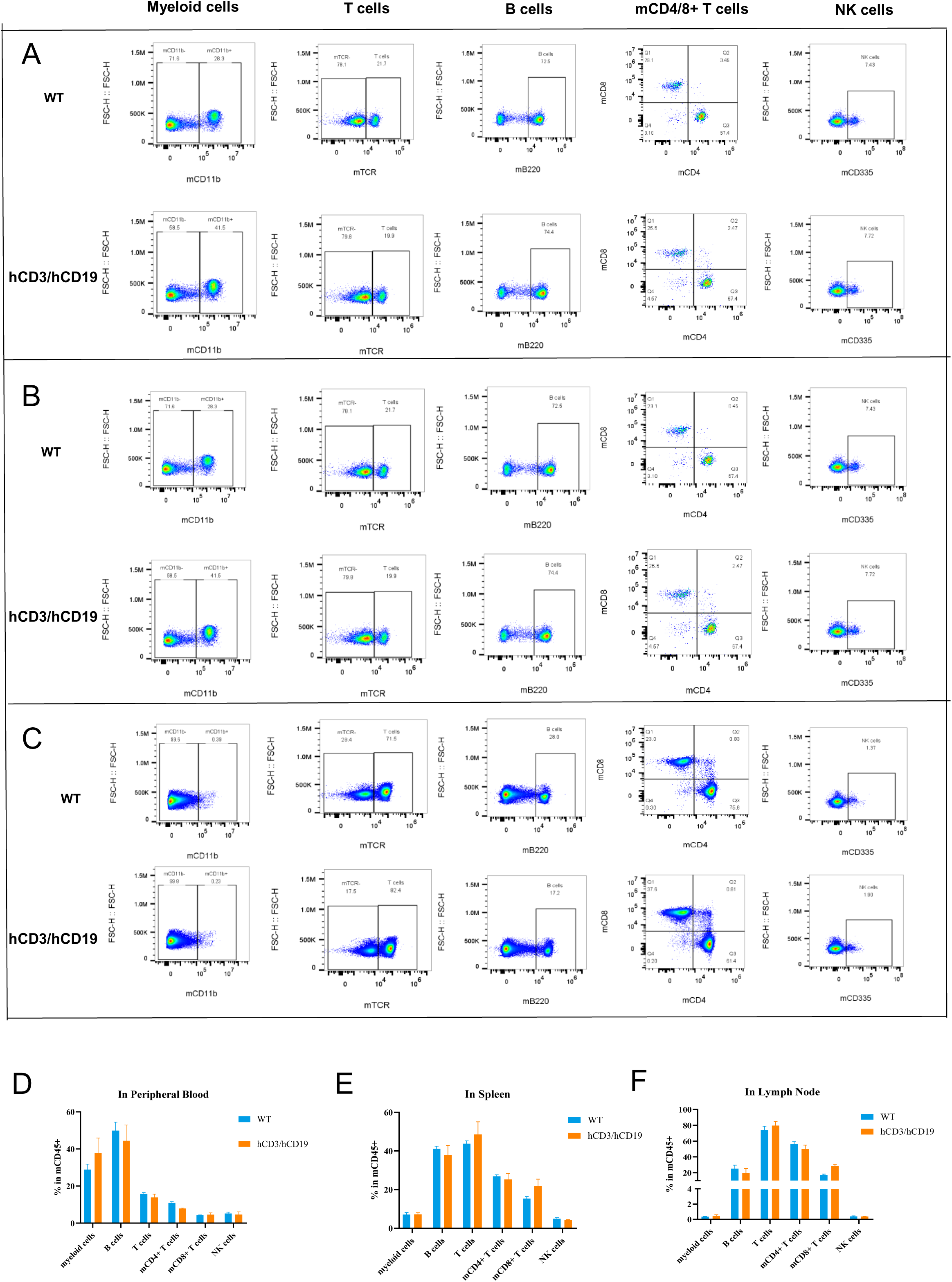
Analysis of immune cell subpopulations in different immune compartments. (A) Representative flow cytometry plots of immune cell subsets in peripheral blood. (B) Immune cell profiles in the spleen by flow cytometry. (C) Immune cell profiles in lymph nodes by flow cytometry. (D–F) Distribution of immune cell subtypes in peripheral blood, spleen, and lymph nodes, respectively.

### 3.4 Evaluation of Small Molecule Drug Efficacy in the Modified SLE Model Using hCD3/hCD19 Mice

The modified induction protocol was employed to establish an SLE model in hCD3/hCD19 mice, and the therapeutic efficacy of cyclophosphamide was subsequently assessed (Fig. 4A). Throughout the study period, mice maintained stable body weights within a safe range and exhibited good tolerance to cyclophosphamide treatment (Fig. 4B).

**Fig. 4.**
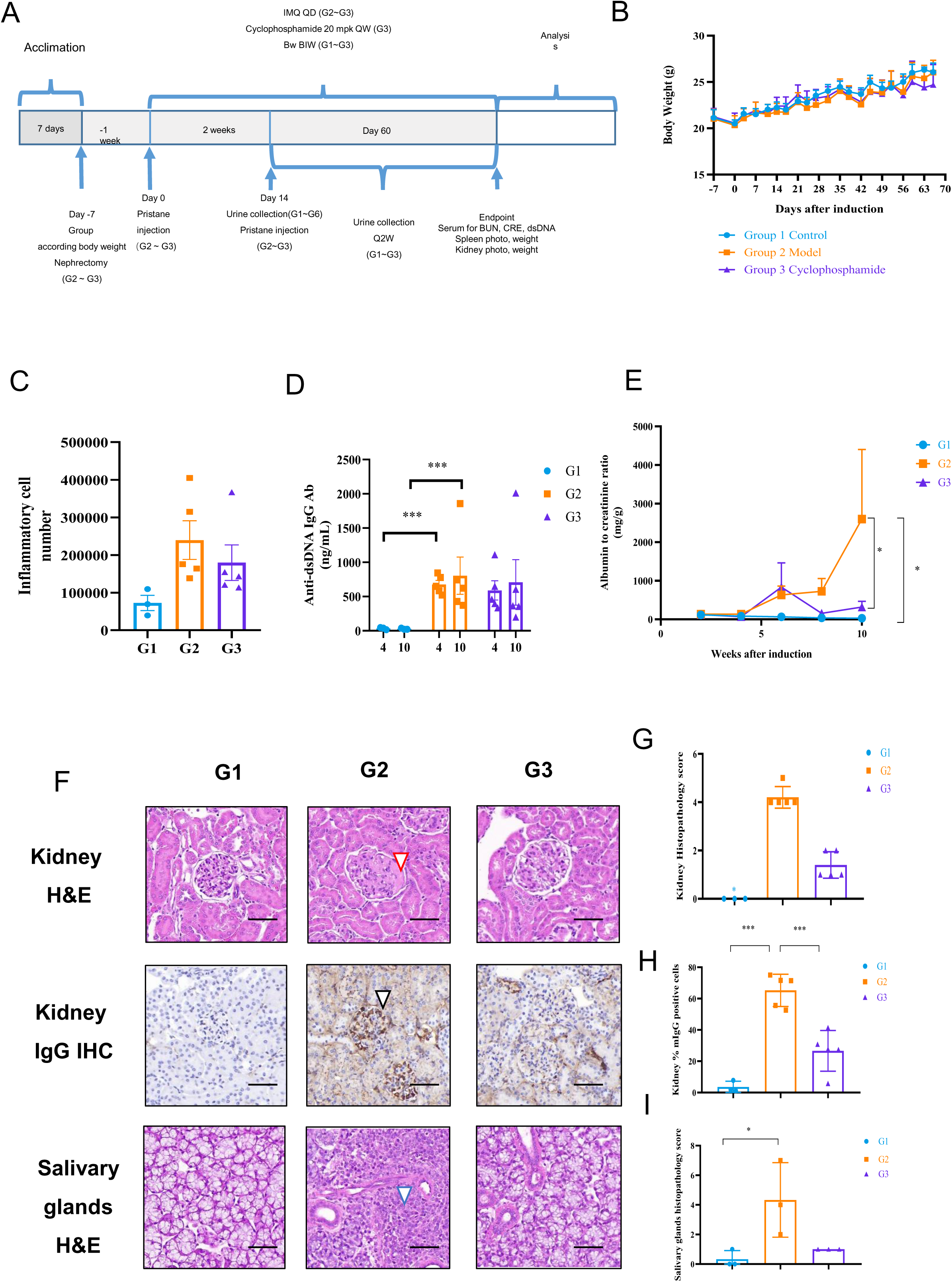
Evaluation of small molecule therapy efficacy in the modified SLE model of hCD3/hCD19 mice. (A) Schematic overview of the modeling and treatment protocol: G1, untreated control; G2, unilateral nephrectomy plus dual Pristane injection and daily IMQ induction; G3, G2 model with cyclophosphamide treatment. (B) Body weight monitoring across the study period for all groups. (C) Quantification of inflammatory cells per 100 µL of peripheral blood at multiple time points. (D) Serum anti-dsDNA IgG levels at weeks 4 and 8. (E) UACR (urinary albumin-to-creatinine ratio) from 24-hour urine collections. (F) Spleen index (spleen weight/body weight) at the endpoint. (G) Representative H&E-stained kidney tissue images from each group. (H) Statistical analysis of kidney damage scores. Sections visualized at 40× magnification; scale bar = 100 μm.

Spleen enlargement was observed in the model mice (Supplemental Fig. 2G), accompanied by a marked increase in circulating inflammatory cells, which was significantly attenuated following cyclophosphamide administration (Fig. 4C). Flow cytometric analysis revealed that myeloid cell populations were elevated, whereas lymphoid cells were reduced; cyclophosphamide treatment effectively reversed these immune cell alterations (Supplemental Fig. 2A–2F).

In addition, serum levels of anti-double-stranded DNA (anti-dsDNA) antibodies were significantly elevated in the model mice and were decreased after cyclophosphamide therapy (Fig. 4D). Urinalysis demonstrated increased urinary albumin excretion and elevated urine albumin-to-creatinine ratio (UACR), both of which were markedly ameliorated by cyclophosphamide treatment (Fig. 4E).

Histological evaluation of renal tissues revealed significant mesangial cell proliferation, glomerular sclerosis, and IgG deposition in the kidneys of model mice, all of which were substantially improved upon cyclophosphamide intervention (Fig. 4F, 4G, and 4H). Moreover, severe inflammatory infiltration was evident in the salivary glands, which was also mitigated following cyclophosphamide treatment (Fig. 4F, 4G, and 4I).

### 3.5 Evaluation of Blincyto Therapeutic Efficacy in the hCD3/hCD19 Mouse SLE Model

The modified SLE model was established in hCD3/hCD19 mice, and the therapeutic potential of Blincyto was evaluated (Fig. 5A). During treatment, body weight fluctuations remained within a safe and manageable range, indicating good tolerance to Blincyto (Fig. 5B).

**Fig. 5.**
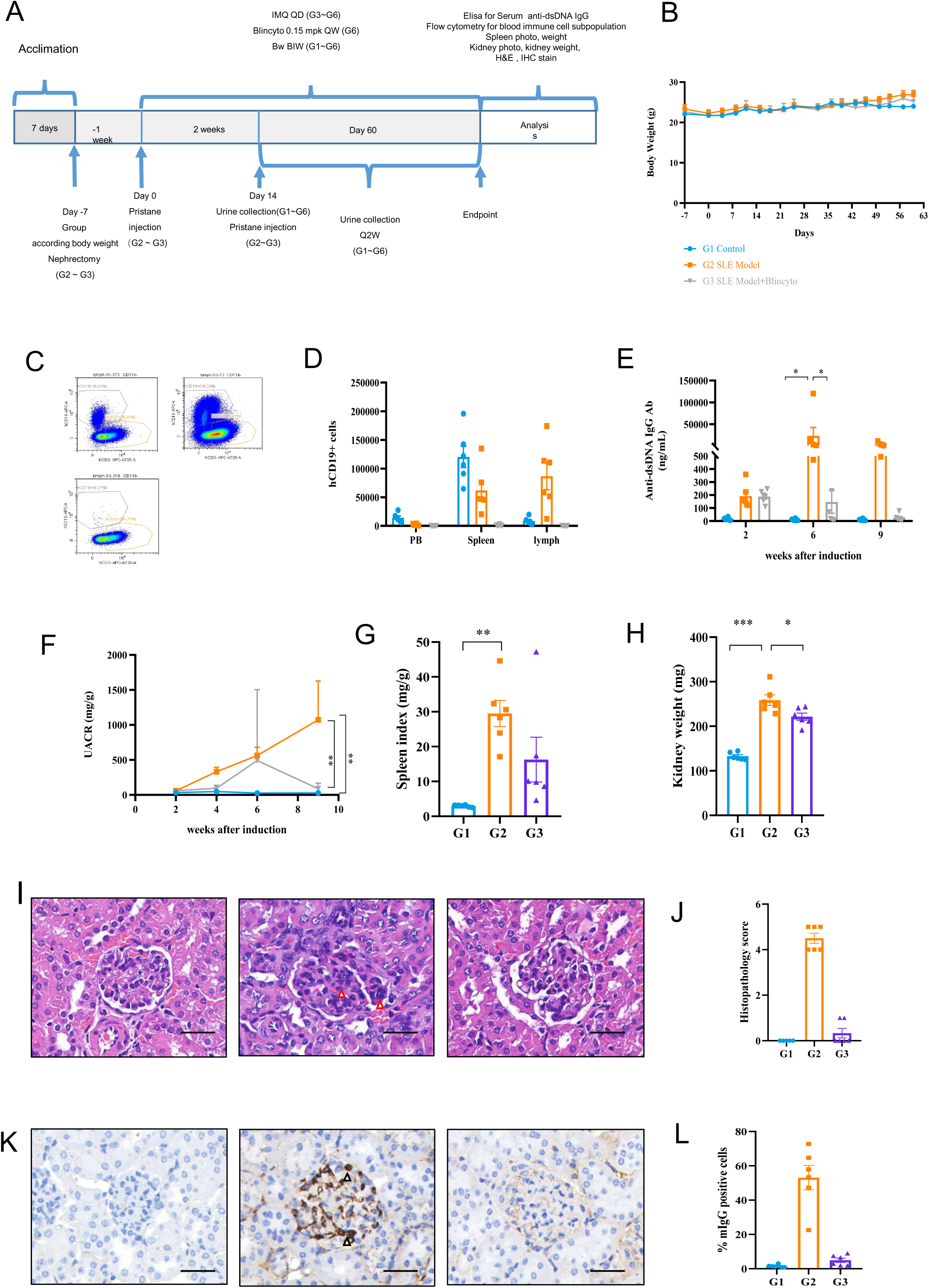
Therapeutic efficacy of Blincyto in the humanized hCD3/hCD19 SLE mouse model. (A) Schematic diagram of the treatment protocol. (B) Body weight trends for each treatment group during the experiment. (C) Representative flow cytometry plot of B cells in lymph nodes. (D) Quantification of B cells in peripheral blood, spleen, and lymph nodes. (E) Serum anti-dsDNA IgG concentrations at weeks 2, 6, and 9. (F) UACR from 24-hour urine samples. (G) Spleen index (spleen weight/body weight) at study termination. (H) Kidney index (kidney weight/body weight) at the endpoint. (I) Representative H&E-stained kidney sections. (J) Summary of kidney pathology scoring. (K) Representative immunohistochemical (IHC) staining of kidneys. (L) Quantification of renal IgG deposition. All sections imaged at 40× magnification; scale bar = 100 μm.

Flow cytometric analysis showed a significant accumulation of B cells in the lymph nodes of model mice, which was markedly reduced following Blincyto administration (Fig. 5C and 5D). Serum anti-double-stranded DNA (anti-dsDNA) antibody levels were substantially elevated in the disease model, and Blincyto treatment significantly lowered these titers (Fig. 5E).

Urinalysis revealed pronounced albuminuria and an elevated urine albumin-to-creatinine ratio (UACR) in the model group, both of which were significantly improved with Blincyto treatment (Fig. 5F). Additionally, spleen enlargement was evident in the model mice, and Blincyto effectively alleviated the hypertrophy of both the spleen and kidneys (Fig. 5G and 5H).

Histopathological examination of the kidneys revealed notable mesangial proliferation and glomerular sclerosis in model mice, which were substantially ameliorated following Blincyto intervention (Fig. 5I and 5J). Furthermore, significant glomerular IgG deposition was observed in the model group and was effectively diminished by Blincyto treatment (Fig. 5K and 5L).

## 4 DISCUSSIONS

The conventional Pristane-induced SLE model is associated with an extended latency period, typically requiring 6–8 months for disease development, alongside only mild renal dysfunction and minimal pathological alterations. Similarly, the IMQ-induced SLE model also presents several limitations. In this study, we optimized the traditional approach by performing unilateral nephrectomy, administering two Pristane injections, and combining this with IMQ induction. Our findings demonstrated that this combined strategy led to a significant elevation in peripheral blood inflammatory cells. At 4- and 8-weeks post-induction, the dsDNA antibody levels were predominantly influenced by IMQ treatment. Moreover, the combined induction exacerbated renal dysfunction, promoted more extensive IgG deposition, and resulted in more severe renal pathology, with similar pathological changes also noted in other organs such as the salivary glands. By 10 weeks, the spleen index showed a stronger association with IMQ administration. Collectively, these results indicate that we successfully established an SLE mouse model with a reduced latency period and clinical indicators in blood, urine, and renal function that more closely mimic human disease.

Peripheral blood analysis revealed that the increase in inflammatory cells was primarily due to an expansion of myeloid populations. Post-modeling, both T and B cell numbers in peripheral blood declined, with T cells particularly sensitive to IMQ treatment. The reduction in B cell counts was observed across both induction protocols, likely reflecting the migration of T and B cells to draining lymph nodes or inflamed tissues.

Using flow cytometry, we confirmed that T cells in hCD3/hCD19 mice expressed humanized surface markers CD3E, CD3D, and CD3G, while B cells expressed humanized CD19. Importantly, the proliferative capacity of T cells following stimulation and the antibody secretion ability of B cells were preserved. Additionally, the distribution of immune cell subtypes in peripheral blood, lymph nodes, and spleen remained within normal ranges.

Building on these findings, we employed the improved induction method to generate an SLE model in hCD3/hCD19 mice. This model exhibited hallmark features of SLE, including elevated inflammatory cell counts in the blood, increased dsDNA antibody levels, compromised renal function, glomerular damage, IgG deposition, and inflammatory infiltration in the salivary glands. Treatment with the classical small-molecule drug cyclophosphamide, previously validated in human and NZB/W mouse studies, significantly ameliorated renal pathology and improved kidney function indicators in the mice.

Following validation of the SLE model and assessment of cyclophosphamide efficacy, we further evaluated the therapeutic potential of a bispecific antibody. Compared to cyclophosphamide, Blincyto demonstrated a more pronounced ability to eliminate B cells and reduce dsDNA antibody levels across multiple tissues. It also led to greater improvements in renal function and histopathological damage. Furthermore, Blincyto treatment resulted in simultaneous improvements in other immune cell populations within the lymph nodes. These results collectively confirm the successful establishment of a humanized mouse SLE model suitable for drug efficacy evaluation.

## 5 CONCLUSIONS

In this study, we established an innovative SLE induction method in Balb/c wildtype and hCD3/hCD19 mice by integrating unilateral nephrectomy, dual pristane injections, and IMQ treatment. This strategy effectively induced SLE-like manifestations, resulting in a validated preclinical mouse model that offers a valuable platform for the evaluation of novel bispecific antibody therapies targeting B cells in SLE.

## 6 STUDY APPROVAL

Experimental animals were housed in the SPF barrier facility at the Shanghai Model Organisms Center, Inc. and acclimatized for 3-7 days prior to use. All procedures were conducted in accordance with IACUC guidelines (Project license No. 2023-0035) approved by the Experimental Animal Ethics Committee of the Shanghai Model Organisms Center, Inc.

## ACKNOWLEDGMENTS

Thanks ChatGPT for proofread and graphical abstract picture drawing.

---

**Supplemental Fig. 1.** (A–F) Quantification of myeloid cells, B cells, plasma cells, cytotoxic T cells, helper T cells, and Treg cells per 100 µL of peripheral blood across groups. (G) Measurement of albumin concentration in 24-hour urine samples. (H) Measurement of creatinine concentration in 24-hour urine samples. (I) Statistical analysis of IgG deposition in kidneys. (J) Representative H&E-stained sections of salivary glands. (K) Histological scoring of salivary gland inflammation. Salivary gland sections were observed at 200× magnification; scale bar = 200 μm.

**Supplemental Fig. 2.** (A) Gene editing strategy schematic for humanized CD3E, CD3D, and CD3G. (B) Gene editing schematic for humanized CD19. (C–E) Proportions of total T cells, cytotoxic T cells, and helper T cells within the inflammatory cell populations.

**Supplemental Fig. 3.** (A–F) Quantification of myeloid cells, B cells, plasma cells, cytotoxic T cells, helper T cells, and Treg cells per 100 µL of peripheral blood.

**Supplemental Fig. 4.** (A–H) Quantitative analysis of inflammatory cells, myeloid cells, B cells, plasma cells, cytotoxic T cells, helper T cells, and Treg cells in peripheral blood, spleen, and lymph nodes across groups.

