## Supplementary figures and images for "Establishment of a Systemic Lupus Erythematosus Mouse Model in Humanized CD3/CD19 Mice and Evaluation of the Efficacy of a Human Bispecific Antibody"

### s1

#### Supplemental fig 1

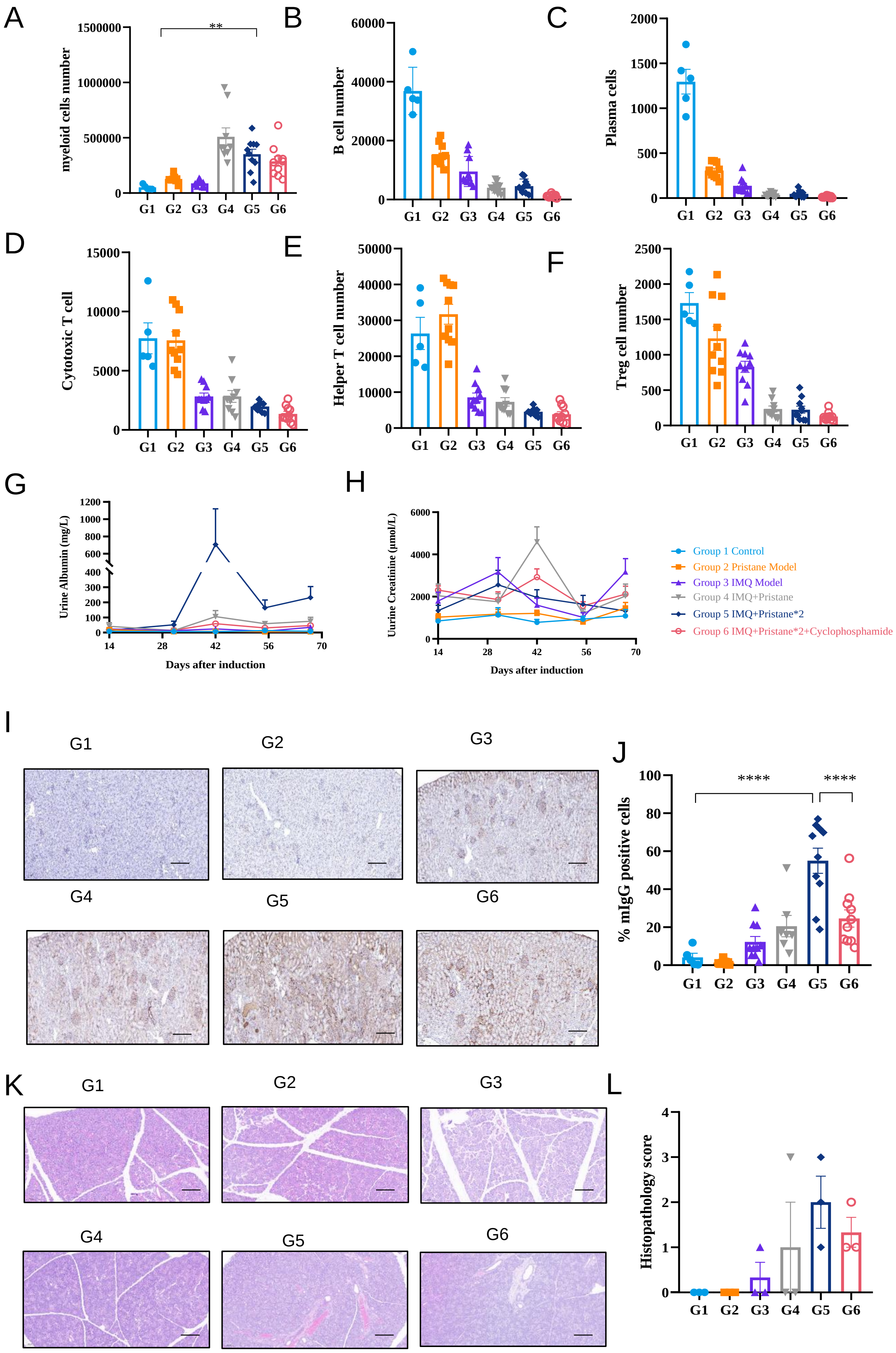

### s2

Supplemental fig 2

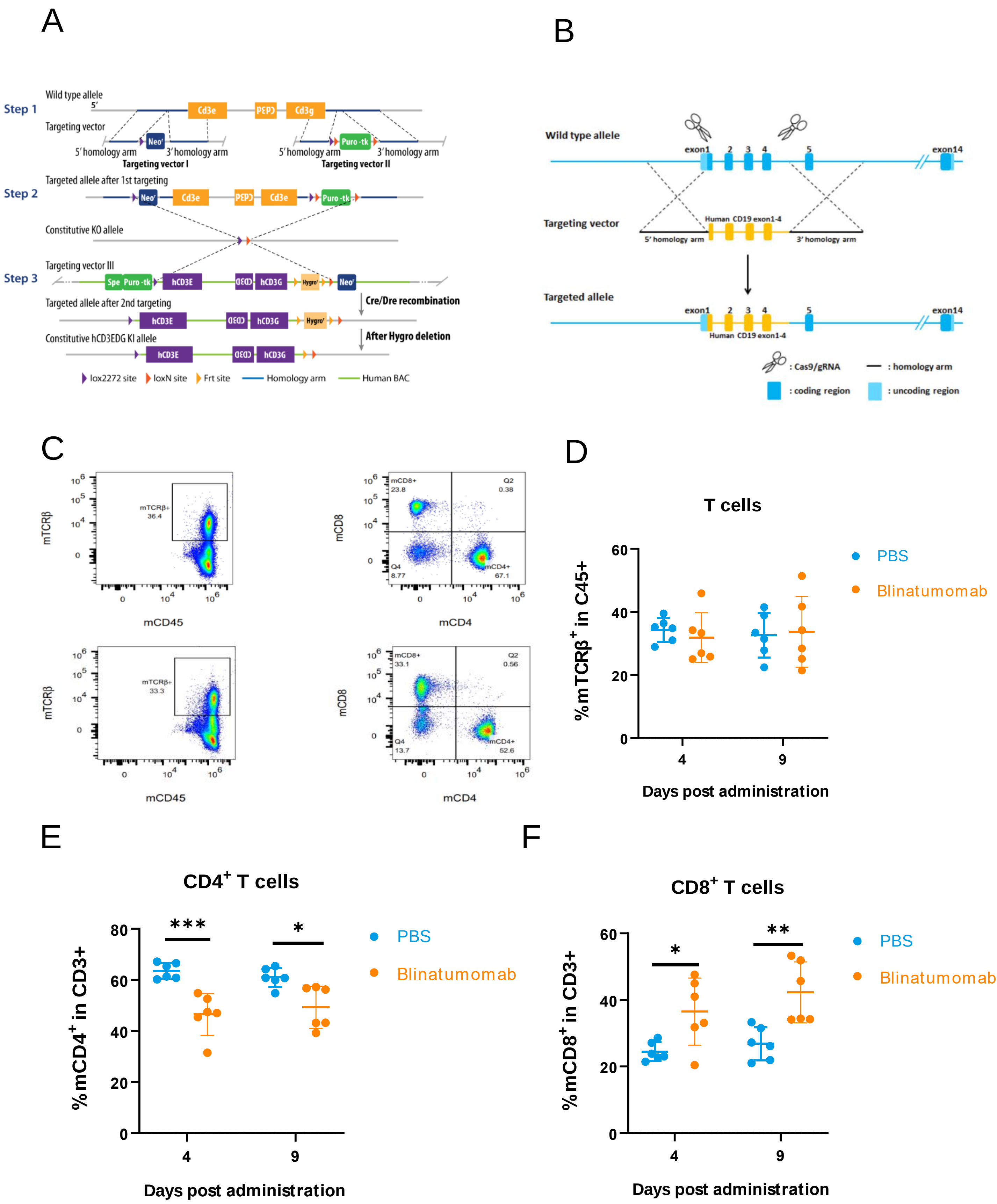

### s3

#### Supplemental fig 3

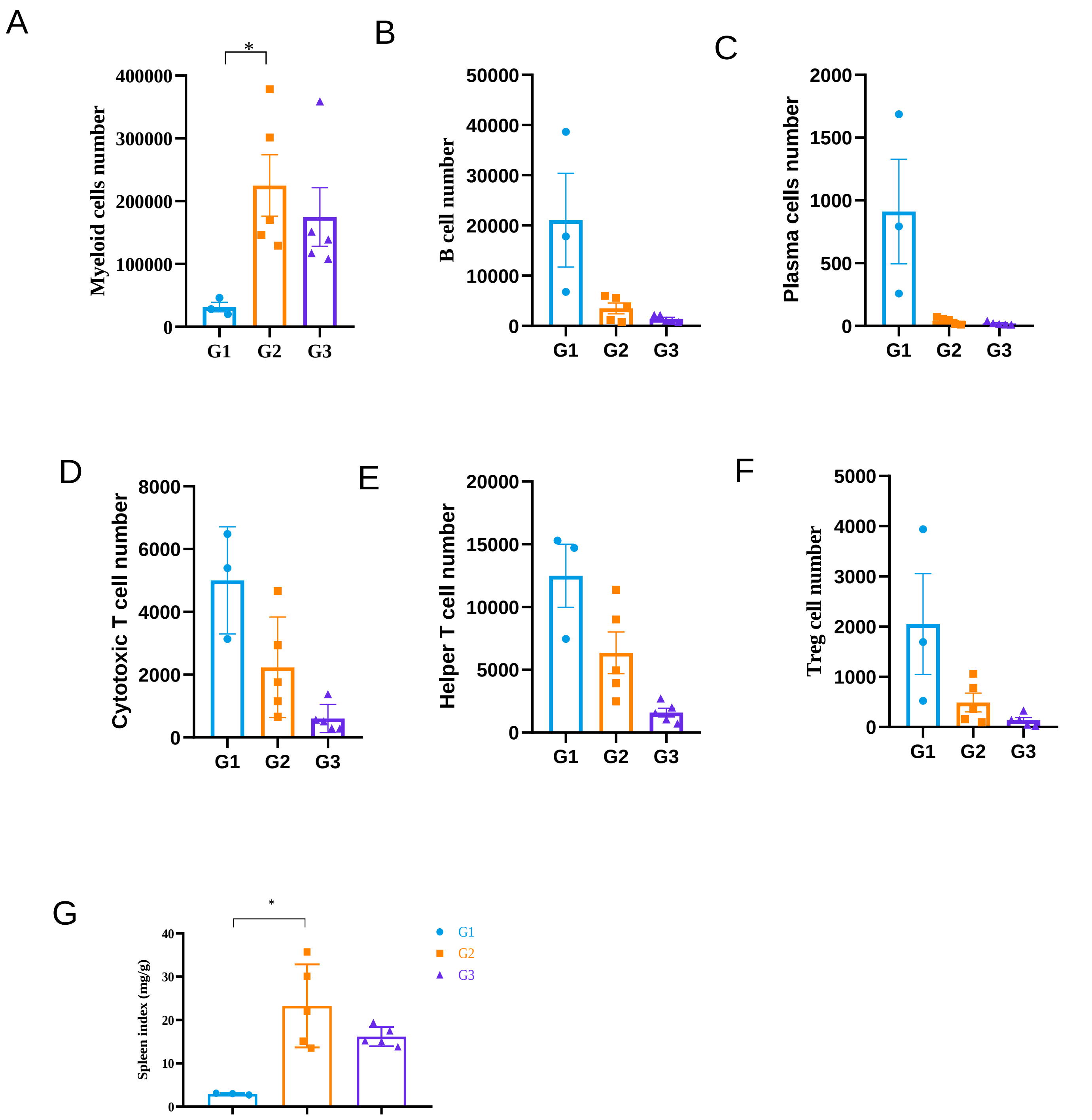

### s4

#### Supplemental fig 4

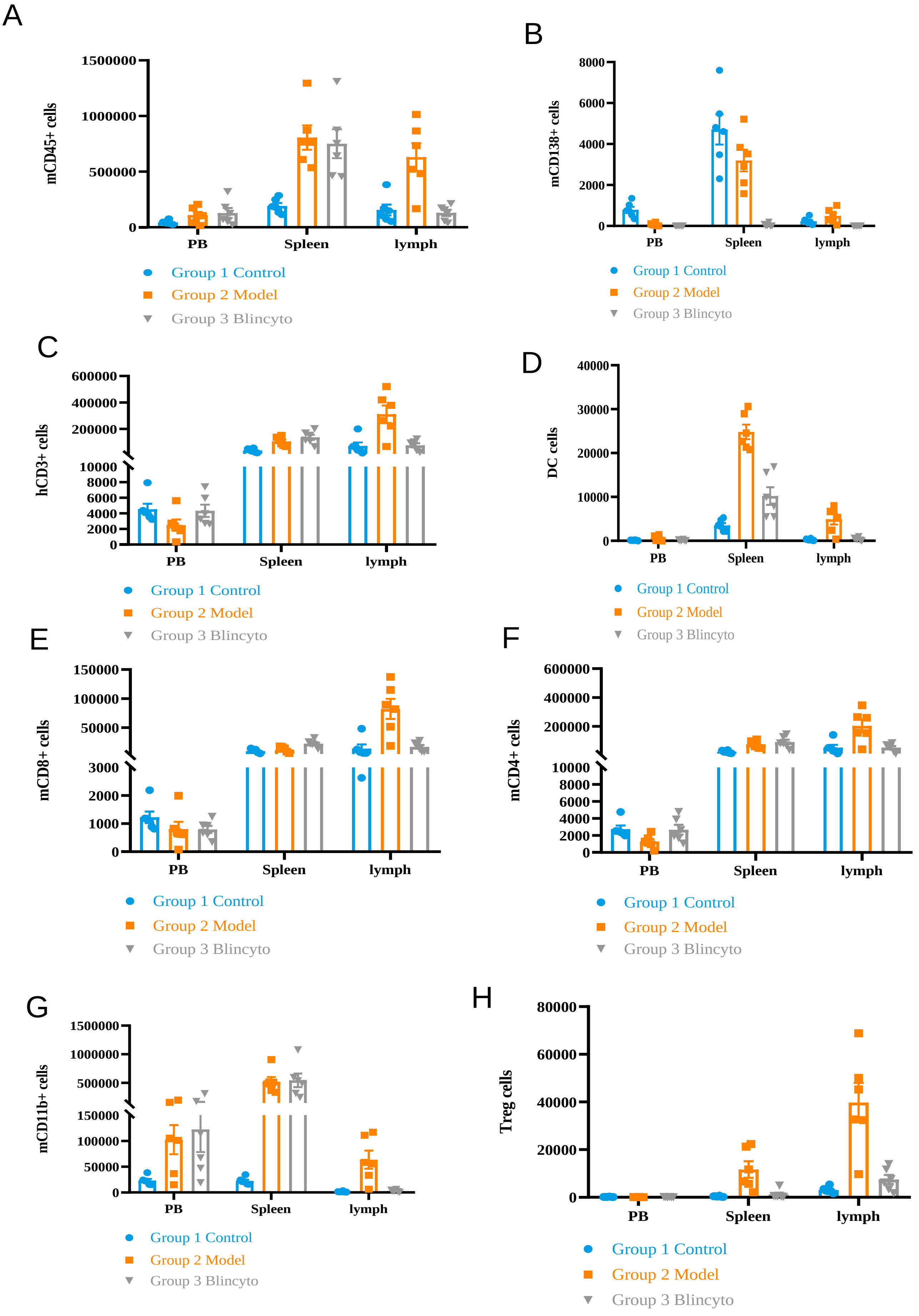
